# Design, synthesis and profiling of highly potent antivirals targeting emerging drug-resistant HIV-1 variants

**DOI:** 10.64898/2025.12.29.696941

**Authors:** Nicolas Jamey, D.M. Nirosh Udayanga, Huanchun Zhang, Shreya M. Ravichandran, Xinyong Cai, Zachary C. Lorson, William M. McFadden, Won Hee Ryu, Roshan Katekar, Jiashu Xie, Karen A. Kirby, Stefan G. Sarafianos, Zhengqiang Wang

## Abstract

Lenacapavir (LEN), the first-in-class HIV capsid inhibitor (CAI), is approved by FDA as a long acting injectable (LAI) for both treatment and pre-exposure prophylaxis (PrEP). Despite its exceptional potency and long pharmacokinetics (PK), a few major resistant mutations have been selected in LEN-treated patients, underscoring the need to develop second-generation LEN analogs to mitigate resistance. Particularly, the M66I mutation confers an extraordinarily high-level LEN resistance, essentially abrogating LEN potency. In this work, we have designed and synthesized LEN analogs featuring a cycloalkyl R^2^ in subunit B drastically different from known analogs. Against wild-type HIV-1, the potency of our analog **3** (EC_50_ = 0.073 nM) was 2.6-fold higher than LEN (EC_50_ = 0.19 nM). More importantly, against the M66I mutant, **3** (EC_50_ = 5.8 nM) was decisively more potent than LEN (EC_50_ > 15 μM) or any known analogs. We have also shown that the size of the R^2^ cycloalkyl ring is a major pharmacophore factor as a smaller (cyclopropyl, analog **1**) or bigger (cyclohexyl, analog **4**) ring confers weaker antiviral potency against both WT HIV-1 and M66I. The vastly improved profile of our lead **3** against M66I was confirmed in the target binding thermal shift assay. These results strongly validate our design and may represent a breakthrough in LEN-based HIV therapy and prophylaxis.

## 1. Introduction

HIV capsid protein (CA) plays a few essential roles in the viral infection cycle: in the early stages, it facilitates cytoplasmic trafficking,^1^ reverse transcription,^2,3^ nuclear import,^4^ uncoating,^5^ and integration targeting,^6^ regulated by various CA-host interactions;^7,8^ in the late stages, newly produced CA assembles into the viral core consisting of approximately 250 hexamers and exactly 12 pentamers,^9,10^ enabled by the protein-protein interactions between CAs.^11^ HIV CA as a drug target^12^ has been clinically validated by the development^13,14^ and FDA-approval of lenacapavir (LEN), a long acting injectable (LAI) administered twice a year. Approved as a therapeutic^15^ and for pre-exposure prophylaxis (PrEP),^16^ LEN could fundamentally alter the paradigm of HIV clinical management and possibly end the pandemic. However, resistance-associated mutations (RAMs) have emerged in LEN-treated populations,^17-19^ of which the major single mutations include Q67H (low fold-resistance, FR), N74D (intermediate FR) and M66I (very high FR).^19^ Mitigating these resistance mutations by designing and synthesizing novel LEN analogs could yield highly coveted second-generation CA-targeting drugs.

In particular, M66I confers complete loss of LEN potency (>80,000 fold-resistance)^20^ and is selected in patients despite the very high fitness price (replication capacity reduced to 4%).^19^ LEN-CA structural analysis^21^ showed that while the M66 in the WT CA interacts comfortably with the di-F phenyl ring (R^2^, subunit B) and the bicyclic ring (R^3^, subunit A) of LEN, I66 creates significant steric clashes via β-branching to severely hamper LEN binding (Figure 1, B). In a recent report,^22^ KFA-027 (Figure 1, C), an R^2^ defluorinated analog (R^2^ = Ph) of LEN (R^2^ = di-F Ph), exhibited slightly moderated (3-fold) potency against WT HIV-1, but substantially better potency than LEN against the M66I mutant (Figure 1, D). Although the improved potency against M66I is desirable, the structural modification is minimal, especially considering that F is a well-known bioisostere for H;^23^ and the sub-μM potency against M66I is modest.

**Figure 1.**
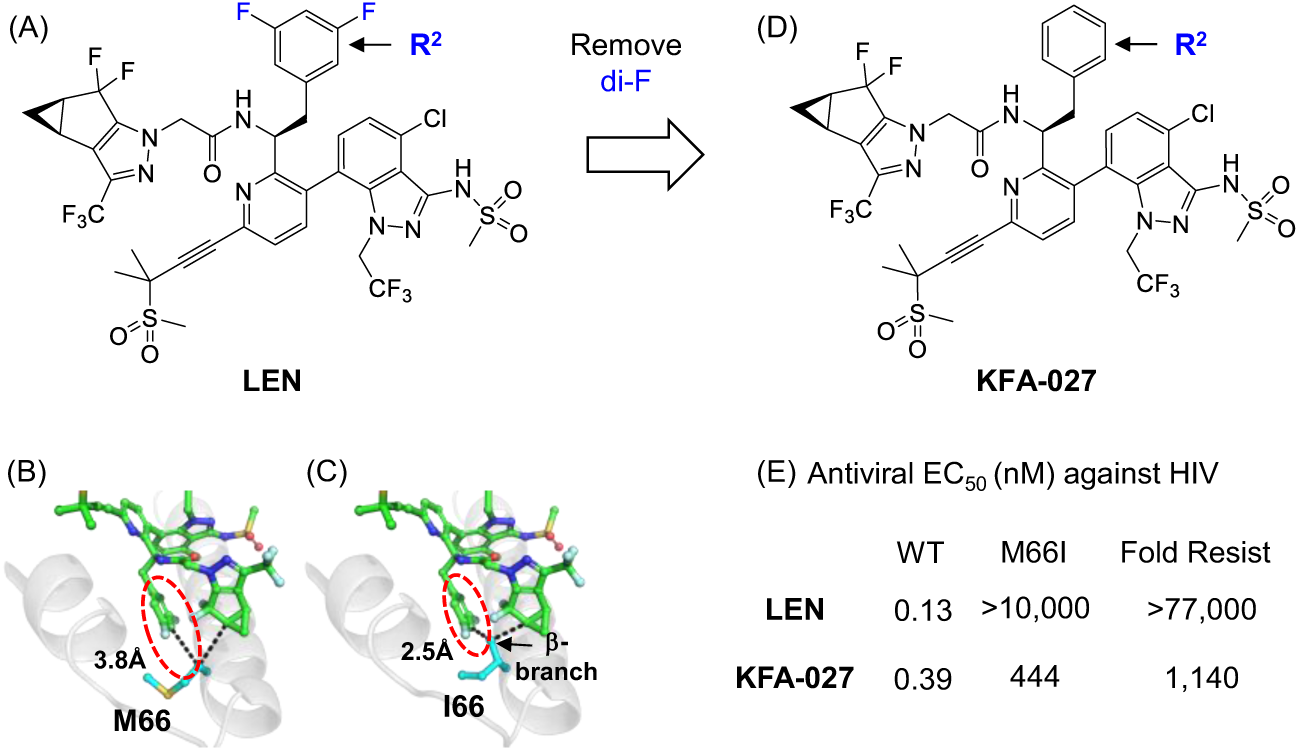
Molecular basis of M66I resistance and its mitigation by reported work. A) Structure of LEN; (B) R^2^ interacts with M66 at an optimal distance; (C) steric hindrance induced by the β-branching of I66; C) structure of R^2^ de-fluorinated analog KFA-027; D) antiviral and resistance profile of LEN and KFA-027.

Concurrent to the reported efforts, we have designed and synthesized subunit B / R^2^ analogs where the di-F phenyl ring of LEN is replaced by a cycloalkyl group (Figure 2). Structurally, these new analogs feature an R^2^ distinct from that of LEN or known analogs in size, electronic and dimensionality. Remarkably, with these structural features, the resulting analog **3** produced overwhelmingly superior antiviral and target binding profiles over LEN against the M66I mutant, while also demonstrating 2.6-fold more potency than LEN against WT HIV-1.

**Figure 2.**
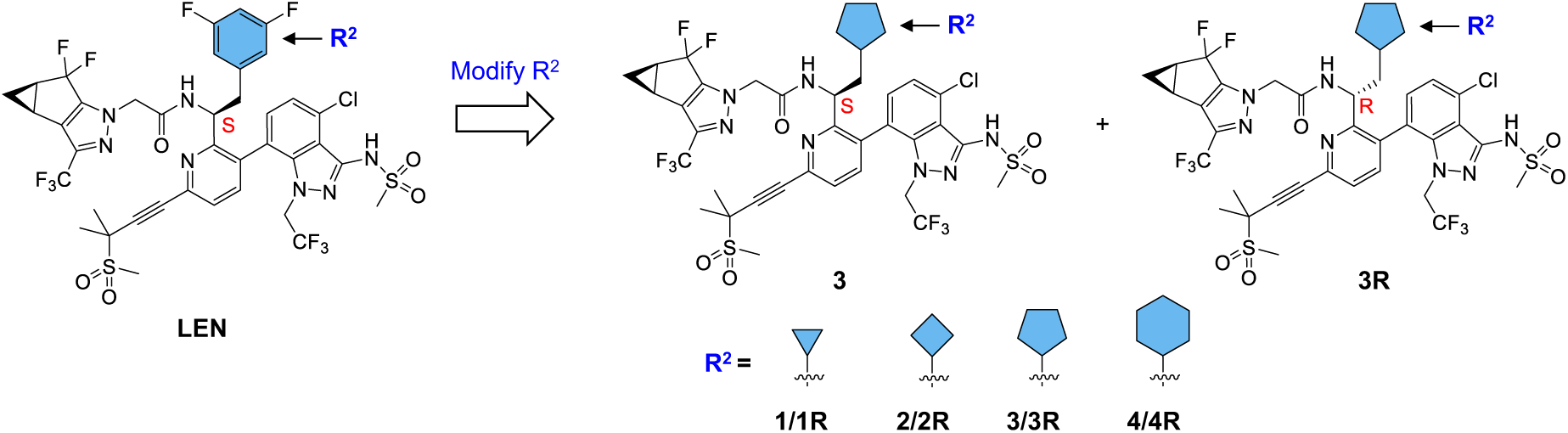
Our designed R^2^ analogs substituting a cycloalkyl for di-F phenyl. Both the size and the nature of the R^2^ group are drastically altered.

## 2. Results and Discussion

### 2.1. Chemical synthesis

The overall synthetic strategy toward our subunit B analogs entails a modular synthesis, combining three subunits named A, B and C (Figure 3). Subunits A and C are the same as used for LEN, whereas structural changes (R group) are pursued within subunit B. The effective synthesis of our subunit B analogs requires developing a convenient procedure described later, starting from 2,5-dibromopyridine instead of 3,6-dibromo-2-methylpyridine. For the subunit A, while reported procedures used a Simmons-Smith cyclopropanation starting with 3-cyclopentenol,^20,24^ the same preparation is not efficient for our scale-up synthesis and does not allow control of the stereochemistry of the newly formed cyclopropane ring. Thus, a new procedure starting with (R)-epichlorohydrin is used in our synthesis as described below. Subunit C is synthesized based on reported procedures.^20^

**Figure 3.**
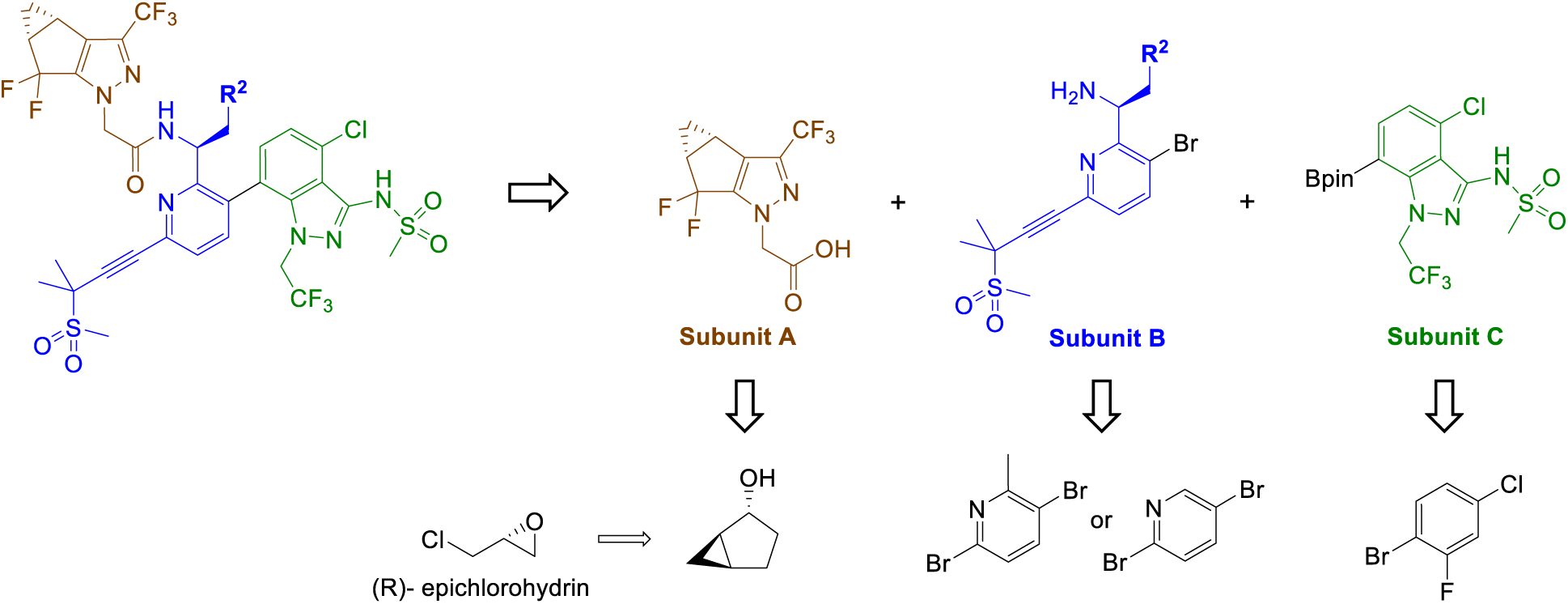
Retrosynthesis of LEN analogs based on a modular synthesis.

Our synthesis of subunit A followed a procedure described by N. Kadunce et al,^25^ for all steps except d, e, and l (Scheme 1). This procedure uses the stereochemistry of the (R)- epichlorohydrin (**5**) to obtain the enantiomerically pure subunit A. In the real synthetic event, (R)- epichlorohydrin (**5**) reacted with allylmagnesium chloride to give epoxide **12c** under basic conditions via alcohol **6**. The subsequent use of lithium 2,2,6,6-tetramethylpiperidide (LTMP) generated *in-situ* induces an intramolecular cyclopropanation of the terminal epoxide of **7**,^26,27^ providing compound **8**. Oxidation of **8** with DMP led to **9**, which was then transformed into **10** using isoamyl nitrite and TMSCl. At this step, the enantiopurity was verified by chiral HPLC, where only one peak was observed on the chromatogram, with a 99% purity. The carbonyl was then protected as a dithiane **11**. The oxime group was then reconverted into carbonyl **12**, which was deprotonated to react with ethyl trifluoroacetate (ETFA), affording 1,3-dione **13**. This 1,3-dione was converted into salt **14** with diisopropylamine. Then the pyrazole ring on **15** was formed by reacting **14** with hydrazinoacetate hydrochloride. The enantiopurity was once again checked at this step by proton NMR with Eu(tfc)_3_ as chiral lanthanide shift reagents,^28^ and as expected no duplication of signals was observed. The difluorination of the dithiane of **15** gave **16**, which upon saponification produced subunit A (Scheme 1).

**Scheme 1.**
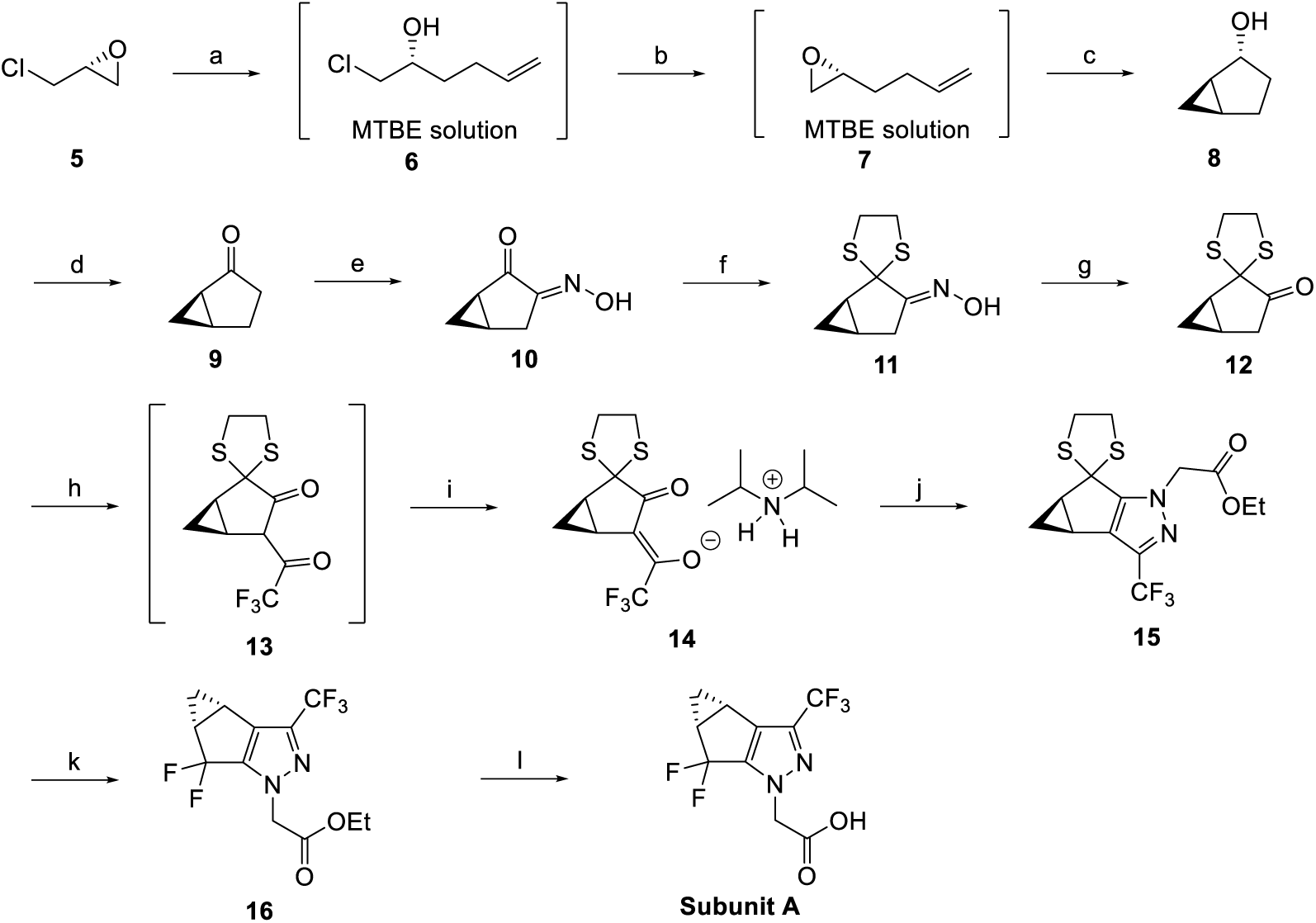
Synthesis of Subunit A*^a^* *^a^*Reagents and conditions: (a) Allylmagnesium chloride, THF, -30 °C, 3h; (b) KOH, MTBE, 12h, 47% over 2 steps; (c) *n*-BuLi, 2,2,6,6-TMP, MTBE, 0 °C, 4h, 96%; (d) DMP, DCM, 0 °C to rt, 12h, 90%; (e) isoamyl nitrite, TMSCl, DCM, -30 °C, 4h, 27%; (f) 1,2-Ethanedithiol, pTSOH, HOAc, rt, overnight, 82%; (g), pTSOH, 2-butanone, water, 75°C, overnight, 89%; (h) LiHMDS, ETFA, THF, -10 °C, 4h; (i) DIPA, rt, 66% over 2 steps; (j) Ethyl hydrazinoacetate hydrochloride, LiCl, CH_3_COCl, EtOH, 55 °C, 18h, 78%; (k) DBDMH, HF-Pyridine, DCM, -15 °C, 80%; (l) LiOH, THF/MeOH, rt, 1h, 95%.

The synthesis of subunit B featuring a cyclopentyl group commenced with commercially available 3,6-dibromo-2-methylpyridine **17** (Scheme 2).^14,20,29^ Bromination of the methyl group on the pyridine ring with *N*-bromosuccinimide and azobisisobutyronitrile furnished geminal dihalide intermediate **18**, which, upon hydrolysis, yielded aldehyde **19**. Condensation of aldehyde **19** with (*S*)-(-)-*tert*-butanesulfinamide afforded *N*-sulfinyl imine **20**. Diastereoselective nucleophilic addition^30^ of cyclopentylmethylmagnesium bromide **21** to imine **20** provided two separable diastereomers (**22**/**22R**). Each diastereomer was subsequently deprotected under acidic conditions to give the corresponding ammonium salt, which was then treated with di-*tert*-butyl dicarbonate and triethylamine to afford enantiopure *N*-Boc-protected bis-bromopyridine intermediates (**23**/**23R**). Finally, Sonogashira cross-coupling of **23**/**23R** with alkyne **28** in the presence of a palladium catalyst, copper(I) iodide, and base delivered the target subunits B **24**/**24R**.

**Scheme 2.**
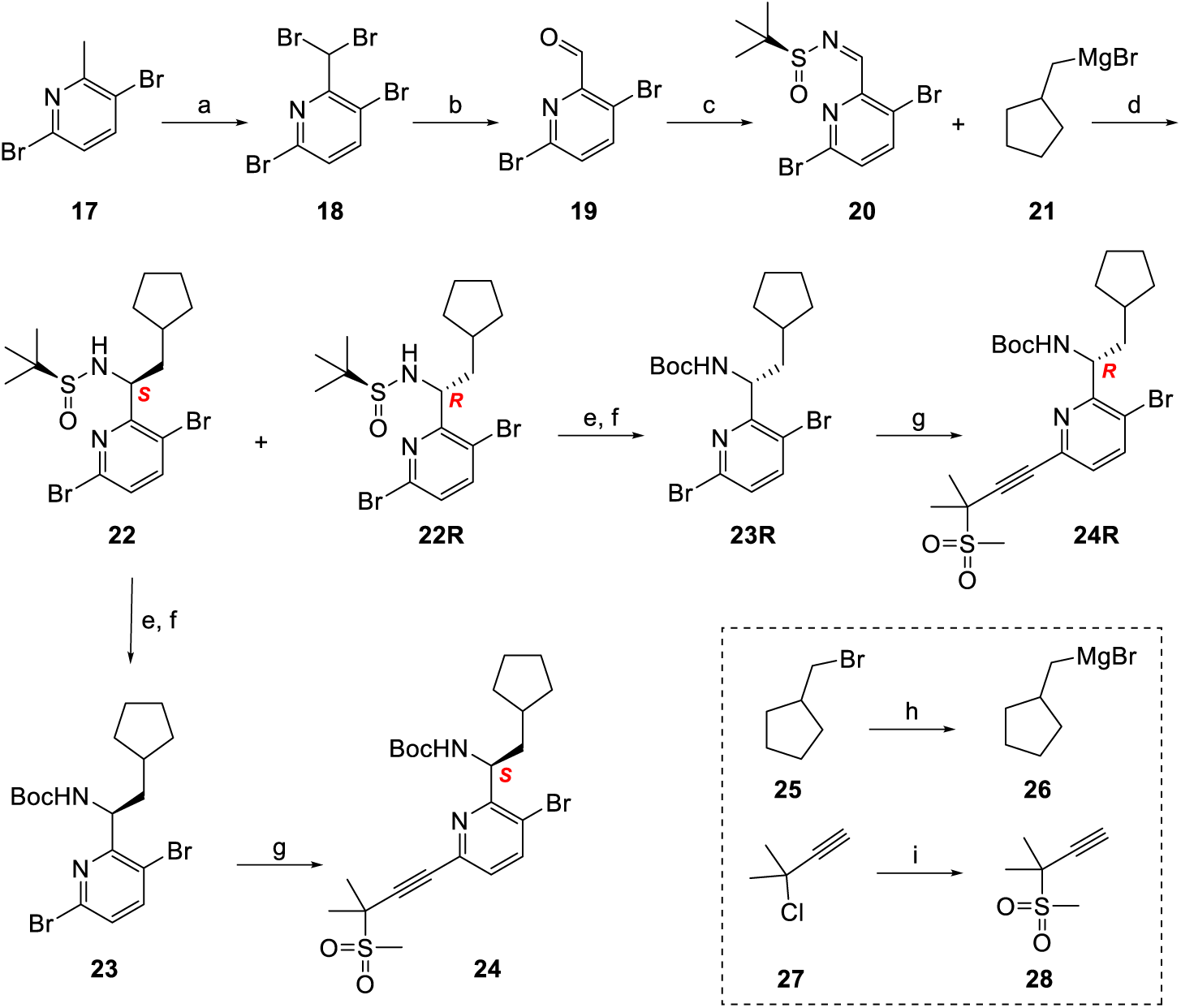
Synthesis of cyclopentyl Subunit B*^a^* *^a^*Reagents and conditions: (a) NBS, AIBN, CCl_4_, 80 °C, 12 h; (b) AgNO_3_, EtOH, H_2_O, reflux, 5 h, 49% (two steps); (c) (*S*)-(-)-tert-Butanesulfinamide, Cs_2_CO_3_, DCM, rt, 1 h, 77%; (d) THF, -78 °C, 2 h; (e) 4 M HCl (in dioxane), MeOH, rt, 3 h; (f) Boc_2_O, Et_3_N, DCM, rt, 2 h, 93-96% (two steps); (g) **28**, Pd(PPh_3_)_2_Cl_2_, CuI, Et_3_N, DMF, rt, 12 h, 50-82%; (h) Mg, I_2_, Et_2_O, rt to 35 °C, 1.5 h; (i) sodium methanesulfinate, Cu(OAc)_2_, TMEDA, iPrOAc, 20 °C to 40 °C, 12 h, 58%.

Synthesis of the cyclopropyl subunit B via the aforementioned procedure was unsuccessful due to cyclopropyl ring opening during the Grignard formation.^31,32^ Consequently, an alternative synthetic route was adopted (Scheme 3),^33-35^ which worked for the synthesis of all cycloalkyl analogs.

**Scheme 3.**
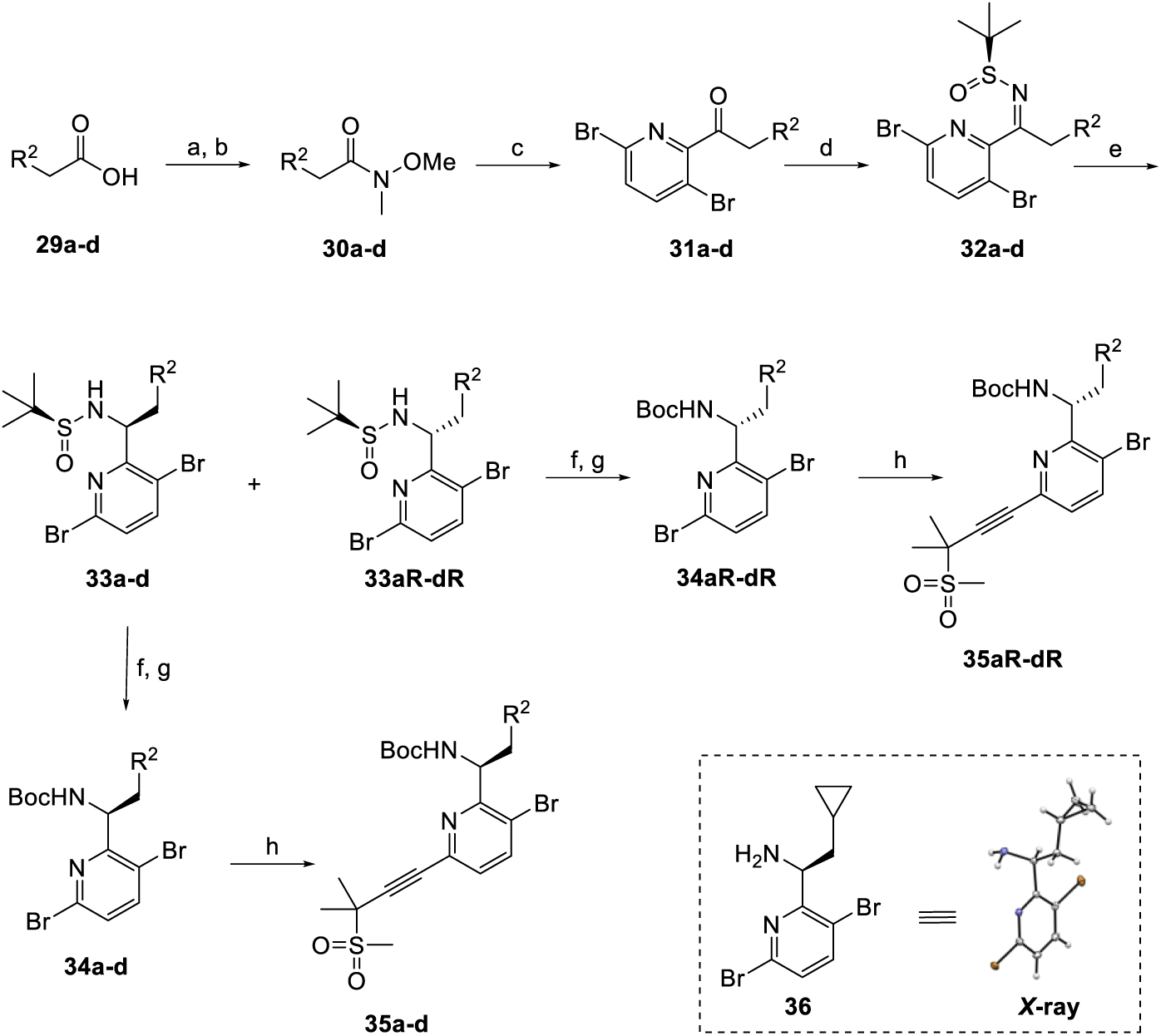
Alternative synthesis of cycloalkyl Subunits B*^a^* *^a^*Reagents and conditions: (a) SOCl_2_, DMF, DCM, rt, 2 h; (b) MeNHOMe.HCl, Et_3_N, DCM, 0 °C to rt, 1 h, 70-85% (two steps); (c) 2,5-dibromopyridine, TMPMgCl.LiCl, THF, -20 °C, 2 h, 70-81%; (d) (*S*)-(-)-tert-butanesulfinamide, Ti(OEt)_4_, THF, 70 °C, 24 h, 47-50%; (e) NaBH_4_, THF, 0 °C to rt, 12 h; (f) 4 M HCl (in dioxane), MeOH, rt, 3 h; (g) Boc_2_O, Et_3_N, DCM, rt, 2 h, 87-95% (two steps); (h) **28**, Pd(PPh_3_)_2_Cl_2_, CuI, Et_3_N, DMF, rt, 12 h, 67-84%.

The synthesis began with commercially available carboxylic acids **29a**−**d**, which were converted to the corresponding Weinreb amides **30a**−**d**. Treatment of Weinreb amides **30a**−**d** with 2,5-dibromopyridine in the presence of TMPMgCl.LiCl afforded ketones **31a**−**d**. Condensation of ketones **31a**−**d** with Ellman’s auxiliary in the presence of titanium(IV) ethoxide furnished *N*-sulfinyl imines **32a**−**d**. Diastereoselective reduction of the imine with sodium borohydride ^36,37^ in THF provided the corresponding diastereomers **33a**/**33aR**−**33d**/**33dR**. Single-crystal *X*-ray crystallographic analysis of the hydrolysis product **36** derived from **34a** confirmed the *S* configuration at the stereogenic center. Removal of the chiral auxiliary under acidic conditions followed by protection with di-*tert*-butyl dicarbonate afforded *N*-Boc-protected amines **34a**/**34aR**−**34d**/**34dR**. Finally, Sonogashira cross-coupling with the methylsulfone-bearing alkyne **28** delivered the target subunits B **35a**/**35aR**−**35d**/**35dR**.

The synthesis of subunit C from 1-bromo-4-chloro-2-fluorobenzene follows a procedure described by F. J. Asturias and M. Kvaratskhelia (Scheme 4).^20^ The procedure started with fomylation of **36** to give aldehyde **37**. The aldehyde was converted into nitrile **38**, followed by a regioselective cyclization by reacting with hydrazine hydrate to afford 3-aminoindazole **39**. The subsequent alkylation of N-1 produced compound **40**, which upon a palladium-catalyzed borylation delivered subunit C.

**Scheme 4.**
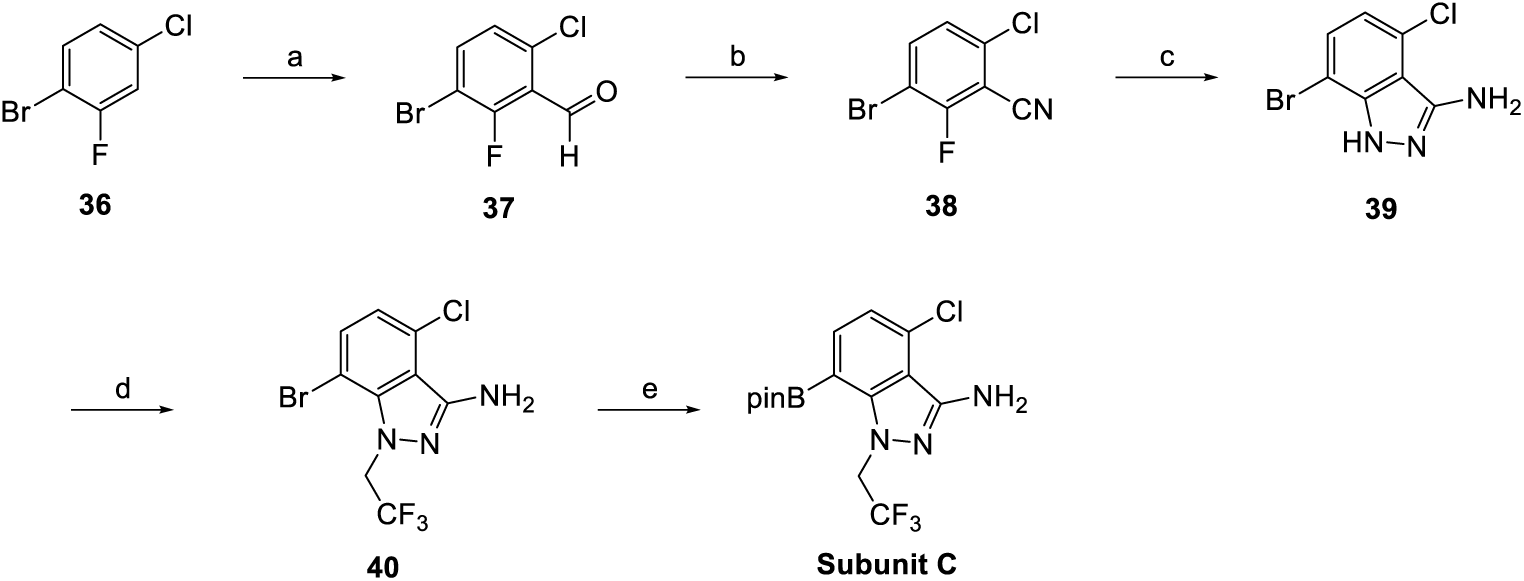
Synthesis of subunit C*^a^* *^a^*Reagents and conditions: (a) LDA, DMF, THF, -78 *°*C, 99%; (b) NH_2_OSO_3_H, H_2_O, 50°C, overnight, 80%; (c) H_2_NNH_2_•H_2_O, EtOH, 80 *°*C, overnight, 27%; (d) 2,2,2-Trifluoroethyl trifluoromethanesulfonate, Cs_2_CO_3_, DMF, 65°C, overnight, 56%; (e) Bpin_2_, Pd(PPh_3_)Cl_2_, KOAc, 1,4-dioxane, 110 *°*C, 70%.

The synthesis of our analogs adopts a reported procedure (Scheme 5).^20^ For the synthesis of (*S*)-isomers, the assembly of subunits B (**35a**−**d**) and C was achieved via Suzuki coupling to afford intermediates **36a**−**d**, which were converted into the sulfonamides **37a**−**d** by addition of methyl sulfonylchloride. From these steps, the presence of two atropoisomers was observed during purification and NMR analysis. Sulfonamides **37a**−**d** were deprotected with TFA, and the amines obtained were directly used in a condensation reaction with the acid of subunit A to give final compounds **1**–**4**. All these steps were also repeated on the *(R)*-enantiomers of subunit B (**35aR**−**dR**) to afford stereoisomeric final compounds **1R**–**4R**. Similar to LEN, each final compound exists as a mixture of two atropoisomers as described below.

**Scheme 5.**
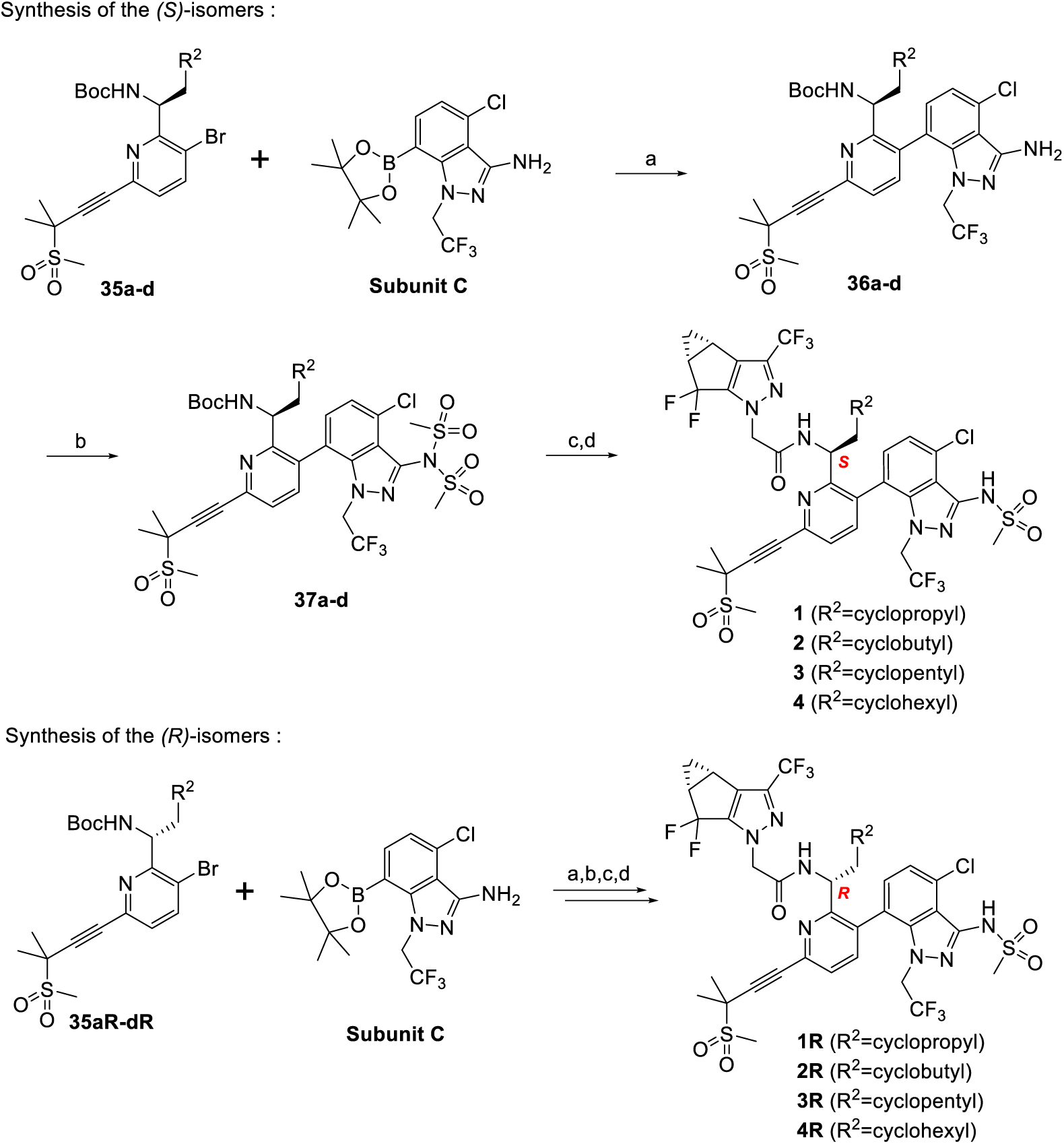
Synthesis of the LEN analogs via subunits B, C and A*^a^* *^a^*Reagents and conditions, all yields with R as cyclopentyl: (a) Pd(dppf)Cl_2_, K_2_CO_3_, 1,4-dioxane/H_2_O, 80 *°*C, 16h, 67%; (b) Methyl sulfonylchloride, Et3N, DCM, rt, 2h, 58%; (c) TFA, DCM, rt, 2h, 99%; (d) Subunit A, HATU, DIPEA, DMF, rt, overnight, 37%.

The two atropoisomers are clearly visible on HPLC and can even be separated. For compound **3** (R^2^=cyclopentyl), the atropisomeric ratio before purification was 1:3.8 (Figure 4). Separation using preparative HPLC yielded a minor atropisomer at 10.784 min and a major one at 11.781 min. However, interconversion was observed with both separated atropoisomers on proton and carbon NMR within hours of separation. Overnight, the separated fraction of the major atropoisomer reached a 0.9:0.1 ratio on ^1^H NMR, and the minor fraction reached a 0.60:0.40 ratio, both favoring the major atropisomer (right peak on HPLC and left peak on 1H NMR, Figure 4).

**Figure 4.**
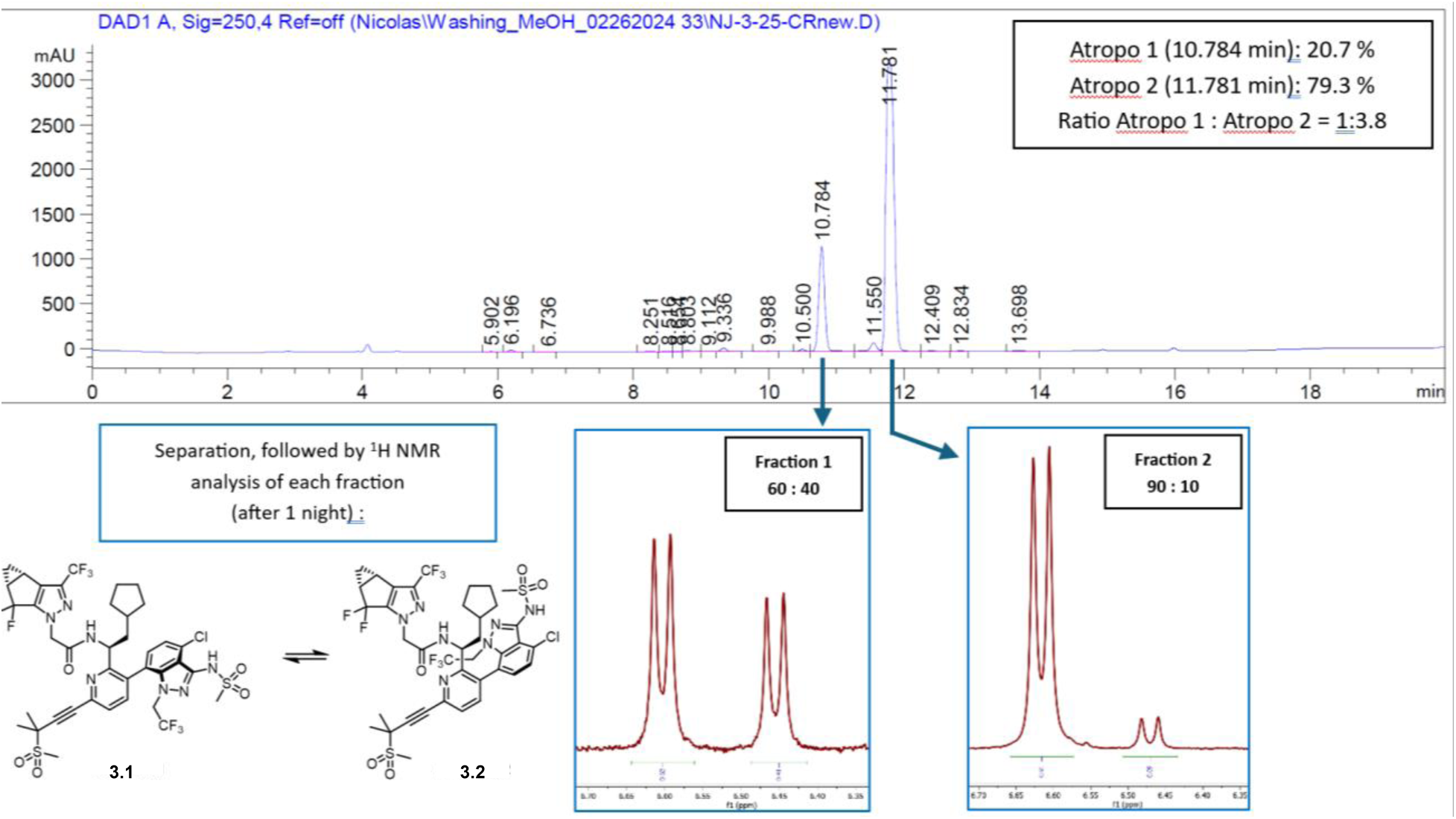
LC-chromatogram of **3** and ^1^H NMR spectra of the two separated atropisomers

These analyses are consistent with the literature on LEN,^20^ which indicates that the atropoisomers are separable only transiently, and interconversion occurs within hours.

### 2.2 Antiviral

New R^2^ analogs were tested against both the WT HIV-1 and the M66I mutant in TZM-GFP reporter cells. LEN was used as both the positive control and the benchmark. Interestingly, of the six analogs tested so far, the three analogs (**1**, **3**, **4**) conforming to the stereochemistry of LEN, all conferred sub to very low nM potencies against WT HIV-1 (Table 1), suggesting that a cycloalkyl group at R^2^ is tolerated. Specifically, in our assay analog **1** (EC_50_ = 0.9 nM) and **4** (EC_50_ = 0.70 nM) were moderately less potent than LEN (EC_50_ = 0.19 nM), whereas analog **3** (EC_50_ = 0.073 nM) produced a potency 2.6-fold higher than LEN. Against the M66I mutant, LEN did not confer significant potency at concentrations up to 15 μM in our assay (Table 1), whereas our analog **4** (EC_50_ = 203 nM) was active in the sub-μM range comparable to the reported potency for KFA-027 (EC_50_ = 444 nM, Figure 1E). Most strikingly, our analogs **1** (EC_50_ = 15.7 nM) and **3** (EC_50_ = 5.8 nM) conferred unprecedented low nM EC_50_ values against M66I (Table 1), drastically more potent than KFA-027^22^ (EC_50_ = 444 nM). The overall superior antiviral profile of our new analogs suggests that the increased *sp*^3^ character^38^ and conformational flexibility^39^ with cycloalkyls largely mitigate the steric clashes while still maintaining excellent target interactions. Our work so far has also revealed two important structure-activity relationship (SAR) trends: 1) the stereochemistry conforming to LEN is required as the R stereoisomers (**1R**, **3R**, **4R**) produced markedly weaker potencies against WT HIV and marginal activities against M66I; 2) analogs with a smaller cyclopropyl (**1**) or bigger cyclohexyl (**4**) are weaker than our lead cyclopentyl analog **3**, suggesting that a 5-membered carbocycle may be the optimal size for R^2^. Collectively, these results strongly validate our analog design.

**Table 1.** Antiviral potency against WT HIV-1 and the M66I mutant.

| Compound | $EC_{50}$ <sup>a</sup> | | FR <sup>d</sup> | CC<sub>50</sub> (μM) |
| --- | --- | --- | --- | --- |
|  | WT HIV-1 (nM) | M66I HIV-1 (nM) |  |  |
| **1** | 0.9 ± 0.4 | 15.7 ± 12.0 | 18 | >100 |
| **1R** | 3.8 ± 0.5 | 2,300 ± 200 | 605 | >100 |
| **3** | 0.073 ± 0.03 | 5.8 ± 4.2 | 79 | >100 |
| **3R** | 12.3 ± 3.5 | 4,000 ± 90 | 325 | >100 |
| **4** | 0.70 ± 0.07 | 203 ± 49 | 290 | >100 |
| **4R** | 70 ± 9 | >40,000 | 570 | >100 |
| **LEN** | 0.19 ± 0.05 <sup>b</sup> | >15,000 <sup>c</sup> | >78,000 | >100 |

<sup>a</sup> In TZM-GFP cells; <sup>b</sup> Published EC<sub>50</sub> = 0.13 nM; <sup>c</sup> Published EC<sub>50</sub> >10 μM; <sup>d</sup> Fold Resistance, defined as EC<sub>50</sub> (M66I) / EC<sub>50</sub> (WT)

### 2.3 Target binding

#### 2.3.1 Thermal shift assay

To further characterize these new analogs, we assessed their direct binding to the target protein in a thermal shift assay (TSA).^40^ Capsid inhibitors are known to bind to CA hexamers but not CA monomers. In this assay, crosslinked CA hexamers of WT, M66I, Q67H, and N74D were used and a larger ΔTm indicates higher target binding affinity. The binding data as shown in Table 2 largely corroborate the observed antiviral potency and SAR trends: 1) in the TSA the three R stereoisomers **1R**, **3R**, and **4R** did not bind at all to any of the tested CA hexamers (Table 2); 2) analogs **1**, **3**, and **4** showed high affinity toward M66I CA hexamer whereas LEN did not bind; 3) **3** produced an overall binding profile superior to that of **1** and **4**. Interestingly, **1**, **3** and **LEN** all bind well to Q67H and N74D CA hexamers at 20 µM compound concentration.

**Table 2.** Thermal shift assay (TSA) measuring compound binding to HIV-1 crosslinked CA hexamers.

| Compound | WT ΔT<sub>m</sub> (°C) | M66I ΔT<sub>m</sub> (°C) | Q67H ΔT<sub>m</sub> (°C) | N74D ΔT<sub>m</sub> (°C) |
| --- | --- | --- | --- | --- |
| **1** | 12.3 ± 0.1 | 9.5 ± 0.1 | 8.0 ± 1.1 | 9.9 ± 0.1 |
| **1R** | 0.07 ± 0.09 | -0.04 ± 0.07 | 1.0 ± 1.1 | 0.1 ± 0.1 |
| **3** | 14.1 ± 0.1 | 10.0 ± 0.2 | 9.6 ± 1.2 | 10.7 ± 0.2 |
| **3R** | 0.0 ± 0.09 | 0.04 ± 0.07 | 1.1 ± 1.1 | 0.1 ± 0.1 |
| **4** | 12.2 ± 0.1 | 7.0 ± 0.1 | NT <sup>a</sup> | NT |
| **4R** | 0.07 ± 0.1 | 0.07 ± 0.1 | NT | NT |
| **LEN** | 16.1 ± 0.4 | -1.1 ± 0.4 | 12.7 ± 1.1 | 13.7 ± 0.1 |

<sup>a</sup> NT = Not Tested

#### 2.3.2 Molecular modeling

To provide computational insights into the difference in activity between LEN and the new analog **3**, we performed induced fit docking (IFD) calculations^41^ generating WT and M66I HIV-1 CA structural models. The starting model for both was 4XFZ, a crystal structure of the WT native full length CA hexamer in complex with PF74.^11^ The existing WT 4XFZ structure was prepared for molecular docking by performing the protein preparation workflow in Schrödinger as described in the Materials and Methods section. The M66I CA structural model was obtained by taking the afore-mentioned prepared 4XFZ structure, performing the M66I mutation on the CA, and performing another round of protein preparation workflow. 4XFZ was used as the reference structure opposed to 6VKV,^20^ of which the cognate ligand is LEN, as the 6VKV structure could not distinguish and explain the difference in activity between LEN and analog **3**.

IFD poses of LEN and analog **3** on the WT CA model show little deviation from the 6VKV crystal structure, maintaining the key protein-ligand interactions (results not shown). IFD on the M66I CA model, however, produces different results. LEN does not dock in the generated M66I CA model with the protein-ligand pose and interactions observed in 6VKV, but rather in a different pose, resulting in the loss of key interactions (Figure 5A). On the other hand, analog **3** docks to CA similar to the observed position in 6VKV, as shown in Figure 5B. Despite the slight difference in how subunit B sits in the binding site, analog **3** maintains the canonical protein-ligand interaction observed in 6VKV. In the absence of LEN-bound structure of CA M66I, the IFD calculations using 4XFZ structure provides a qualitative explanation to the difference in activity between LEN and analog **3**.

**Figure 5.**
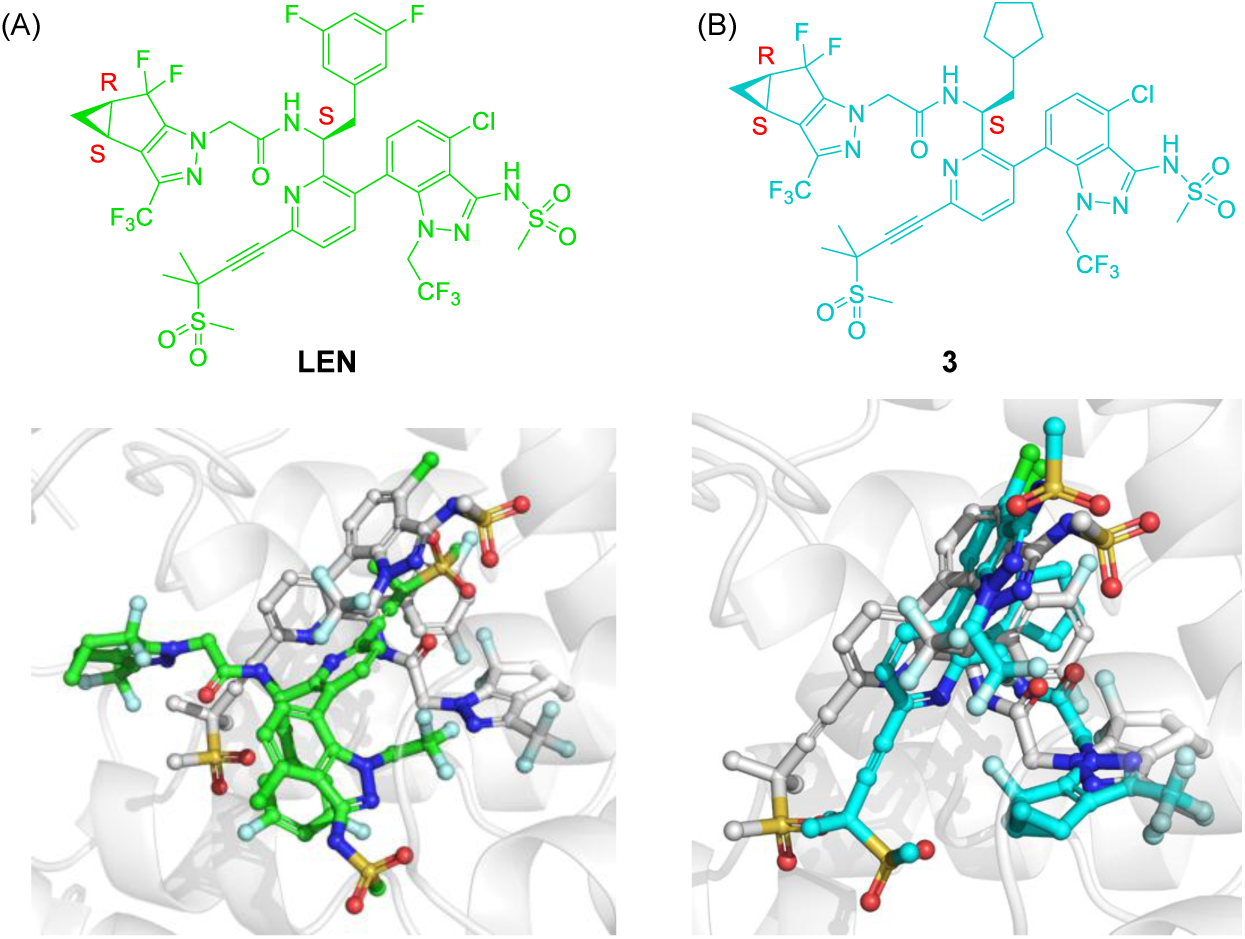
Induced-fit docking (IFD) poses of LEN and analog **3** to a M66I CA structural model. (A) IFD pose of LEN is shown in green and the position of LEN (from PDB: 6VKV) after superposition of the CA proteins is shown in gray. (B) IFD pose of analog **3** is shown in cyan and position of LEN in PDB 6VKV after superposition of the CA proteins is shown in gray.

Our ongoing efforts focus on further exploring the R^2^ SAR and characterizing our strong lead compound **3** for structure and mechanism, ADME profiling, and pharmacokinetics (PK) in rats.

## 3. Conclusion

The M66I mutation within the LEN binding pocket of HIV CA confers complete loss of LEN antiviral potency. Toward mitigating this mutation, previously reported work identified KFA-027 as the LEN analog most potent against M66I to date (EC_50_ = 444 nM). In our work reported herein, we have designed and synthesized novel LEN analogs showing drastically improved potency against the M66I mutant, as exemplified by tested analog **3** (EC_50_ = 5.8 nM). In the thermal shift assay, our analogs produced large shifts against M66I CA hexamer whereas the shift with LEN was marginal, indicating high binding affinity of our analogs and corroborating the exceptional antiviral potency observed. Our analog design features cycloalkyl R^2^ groups radically different from LEN or its known analogs. The decisively superior antiviral potency of our lead **3** over LEN against both M66I and WT HIV-1 strongly validates our analog design strategy. Overall, our novel analogs represent a significant advancement in LEN-based HIV therapy and prophylaxis.

## 4. Materials and Methods

### 4.1. Chemistry

All commercial chemicals were used as supplied unless indicated otherwise. Compounds were purified via flash chromatography using a Combiflash RF-200 (Teledyne ISCO, Lincoln, NE, USA) with RediSep columns (silica) and indicated mobile phase. ^1^H and ^13^C NMR spectra were recorded on a Bruker 400 spectrometer (Bruker, Billerica, MA, USA). Diastereomeric ratio was determined by ^1^H NMR analysis. Mass data was acquired using an Agilent 6230 TOF LC/MS spectrometer (Agilent Technologies). Compound purity analysis was performed using Agilent 1260 Infinity HPLC with an Eclipse C18 column (3.5 μm, 4.6 × 100 mm). HPLC conditions: flow rate, 1.0 mL/min; solvent A, 0.1% TFA in water; solvent B, 0.1% TFA in acetonitrile; gradient (B, %): 0–3 min (5–100), 3–11 min (100), 11–13 min (100–5). Determined purity was >95% for all final compounds.

*N-((S)-1-(3-(4-chloro-3-(methylsulfonamido)-1-(2,2,2-trifluoroethyl)-1H-indazol-7-yl)-6-(3-methyl-3-(methylsulfonyl)but-1-yn-1-yl)pyridin-2-yl)-2-cyclopropylethyl)-2-((3bS,4aR)-5,5-difluoro-3-(trifluoromethyl)-3b,4,4a,5-tetrahydro-1H-cyclopropa[3,4]cyclopenta[1,2-c]pyrazol-1-yl)acetamide* (**1**)

The two atropoisomers of **1** were not separated. Both can be seen in NMR spectra, with a ratio of ∼0.85:0.15.

**^1^H NMR** (400 MHz, CDCl_3_) δ 7.60 (d, *J* = 7.9 Hz, 1H), 7.53 (d, *J* = 11.1 Hz, 1H), 7.51 (d, *J* = 8.0 Hz, 1H), 7.21 (d, *J* = 7.6 Hz, 1H), 7.03 (d, *J* = 7.7 Hz, 1H), 6.86 (d, *J* = 8.4 Hz, 1H), 4.77 (s, 2H), 4.76 – 4.71 (m, 1H), 4.52 (dq, *J* = 16.2, 8.1 Hz, 1H), 3.99 (dq, *J* = 16.1, 8.1 Hz, 1H), 3.41 (s, 3H), 3.16 (s, 3H), 2.57 – 2.42 (m, 2H), 1.83 (s, 6H), 1.59 – 1.48 (m, 1H), 1.47 – 1.37 (m, 2H), 1.17 – 1.09 (m, 1H), 0.42 – 0.32 (m, 1H), 0.31 – 0.25 (m, 1H), 0.26 – 0.15 (m, 1H), -0.12 – -0.21 (m, 1H), -0.39 – -0.47 (m, 1H); **^13^C NMR** (100 MHz, CDCl_3_) δ 164.0, 159.4, 143.6 (t, *J* = 29.3 Hz), 142.7, 139.9, 139.6, 138.9, 136.7 (q, *J* = 39.9 Hz), 132.89 – 132.53 (m), 131.4, 130.0, 126.9, 126.4, 122.8 (q, *J* = 279.6 Hz), 121.7, 120.5 (q, *J* = 267.3 Hz), 119.9 (t, *J* = 243.0 Hz), 119.1, 115.2, 88.1, 85.1, 58.0, 53.7, 51.9, 50.8 (q, *J* = 35.1 Hz), 41.5, 40.9, 35.2, 28.1 (dd, *J* = 34.6, 29.3 Hz), 23.4, 22.7, 12.2, 7.4, 4.7, 4.1; **^19^F NMR** (376 MHz, CDCl_3_) δ -61.57, -70.30, -80.94 (d, *J* = 256.0 Hz), -103.48 (d, *J* = 256.0 Hz) (minor atropisomer not included); **HRMS** (ESI^+^) Calculated for C_36_H_35_ClF_8_N_7_O_5_S_2_ [M+H]^+^ 896.1702; found 896.1695.

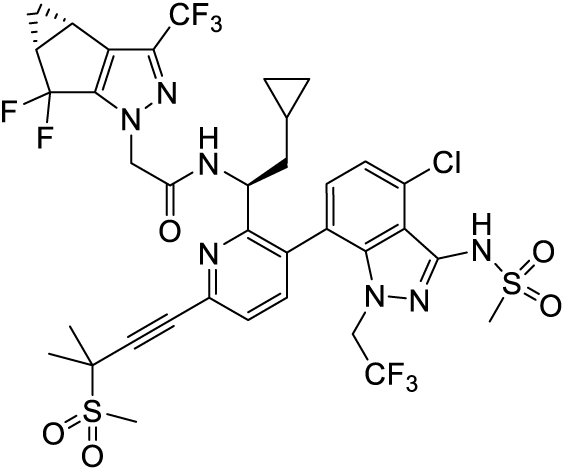

*N-((R)-1-(3-(4-chloro-3-(methylsulfonamido)-1-(2,2,2-trifluoroethyl)-1H-indazol-7-yl)-6-(3-methyl-3-(methylsulfonyl)but-1-yn-1-yl)pyridin-2-yl)-2-cyclopropylethyl)-2-((3bS,4aR)-5,5-difluoro-3-(trifluoromethyl)-3b,4,4a,5-tetrahydro-1H-cyclopropa[3,4]cyclopenta[1,2-c]pyrazol-1-yl)acetamide (**1R**)*.

**^1^H-NMR** (400 MHz, CDCl3) δ ppm: 7.63 (s, 0.15H, minor atropoisomer), 7.58 (d, *J* = 7.9 Hz, 1H), 7.51 – 7.45 (m, 2H), 7.20 (d, *J* = 7.7 Hz, 1H), 7.02 (d, *J* = 7.7 Hz, 1H), 6.72 (d, *J* = 8.5 Hz, 0.85H, major atropoisomer), 6.58 (d, *J* = 8.7 Hz, 0.15H, minor atropoisomer), 5.07 (td, *J* = 8.8, 4.8 Hz, 0.15H), 4.81 – 4.68 (m, 2.9H, major atropoisomer), 4.52 (dq, *J* = 16.2, 8.1 Hz, 0.85H, major atropoisomer), 4.39 (dq, *J* = 15.5, 7.7 Hz, 0.15H, minor atropoisomer), 4.08 (dq, *J* = 16.1, 7.8 Hz, 0.15H, minor atropoisomer), 3.97 (dq, *J* = 16.0, 8.0 Hz, 0.85H, major atropoisomer), 3.41 (2 s, 3H), 3.15 (s, 3H), 2.53 – 2.43 (m, 2H), 1.82 (s, 6H), 1.50 (dq, J = 14.1, 7.0 Hz, 1H), 1.45 – 1.32 (m, 2H), 1.17 – 1.08 (m, 1H), 0.39 – 0.15 (m, 3H), -0.18 (dq, *J* = 9.2, 4.7 Hz, 1H), -0.31 – -0.38 (m, 0.15H, minor atropoisomer), -0.43 (dq, *J* = 9.6, 4.8 Hz, 0.85H, major atropoisomer).**^13^C NMR** (100 MHz, CDCl3) δ 164.1, 159.5, 142.8, 140.1, 139.7, 139.1, 131.5, 130.1, 127.1, 126.5, 121.7, 119.2, 115.2, 88.3, 85.2, 58.1, 53.8, 51.9, 51.9, 41.7, 41.1, 35.3, 28.2 (dd, J = 34.5, 29.4 Hz), 23.5, 22.8, 22.8, 22.73, 12.4, 7.4, 4.8, 4.2; **^19^F NMR** (376 MHz, CDCl_3_) δ -61.53, -70.27, -81.15 (dd, *J* = 255.6, 13.4 Hz), -103.55 (dd, *J* = 256.1, 8.4 Hz).**HRMS** (ESI^+^) Calculated for C_36_H_35_ClF_8_N_7_O_5_S_2_ [M+H]+ 896.1702; found 896.1685.

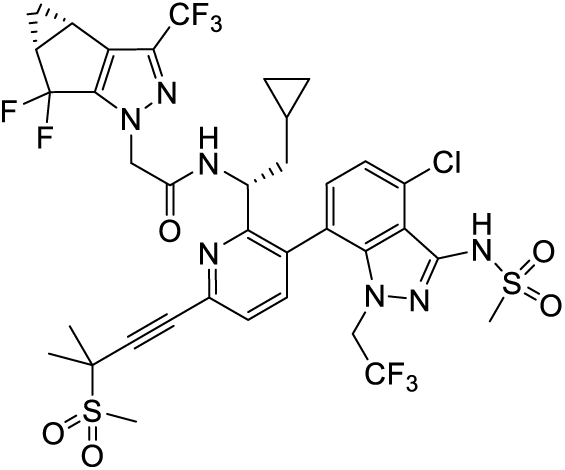

*N-((S)-1-(3-(4-chloro-3-(methylsulfonamido)-1-(2,2,2-trifluoroethyl)-1H-indazol-7-yl)-6-(3-methyl-3-(methylsulfonyl)but-1-yn-1-yl)pyridin-2-yl)-2-cyclobutylethyl)-2-((3bR,4aS)-5,5-difluoro-3-(trifluoromethyl)-3b,4,4a,5-tetrahydro-1H-cyclopropa[3,4]cyclopenta[1,2-c]pyrazol-1-yl)acetamide* (**2**)

**^1^H NMR** (400 MHz, CDCl_3_) δ 7.58 (d, *J* = 8.0 Hz, 1H), 7.48 (d, *J* = 7.9 Hz, 1H), 7.44 (s, 1H), 7.24 (d, *J* = 7.6 Hz, 1H), 7.05 (d, *J* = 7.7 Hz, 1H), 6.65 (d, *J* = 8.7 Hz, 1H), 4.74 (s, 2H), 4.58 – 4.52 (m, 1H), 4.48 (dq, *J* = 16.4, 8.0 Hz, 1H), 3.95 (dq, *J* = 16.0, 8.1 Hz, 1H), 3.40 (s, 3H), 3.15 (s, 3H), 2.56 – 2.43 (m, 2H), 2.03 – 1.93 (m, 1H), 1.83 (s, 6H), 1.79 – 1.60 (m, 5H), 1.58 – 1.50 (m, 1H), 1.47 – 1.39 (m, 1H), 1.37 – 1.29 (m, 1H), 1.17 – 1.08 (m, 1H), 1.06 – 0.94 (m, 1H); **^13^C NMR** (100 MHz, CDCl_3_) δ 163.9, 159.5, 143.5 (t, *J* = 29.0 Hz), 142.6, 139.9, 139.6, 139.0, 136.8 (q, *J* = 39.8 Hz), 132.9 – 132.6 (m), 131.3, 129.6, 127.0, 126.4, 122.7 (q, *J* = 279.8 Hz), 121.7, 120.5 (q, *J* = 267.5 Hz), 119.9 (t, *J* = 242.5 Hz), 118.9, 115.0, 88.2, 85.0, 58.0, 53.7, 50.8 (q, *J* = 34.2 Hz), 50.2, 43.0, 41.6, 35.2, 32.5, 28.3, 28.0 (dd, *J* = 34.3, 29.0 Hz), 27.8, 23.4, 22.7, 22.7, 18.4, 12.3; **^19^F NMR** (376 MHz, CDCl_3_) δ -61.60, -70.34, - 81.03 (d, *J* = 255.9 Hz), -103.50 (d, *J* = 256.0 Hz); (minor atropisomer not included); **HRMS** (ESI^+^) Calculated for C_37_H_37_ClF_8_N_7_O_5_S_2_ [M+H]^+^ 910.1858; found 910.1846.

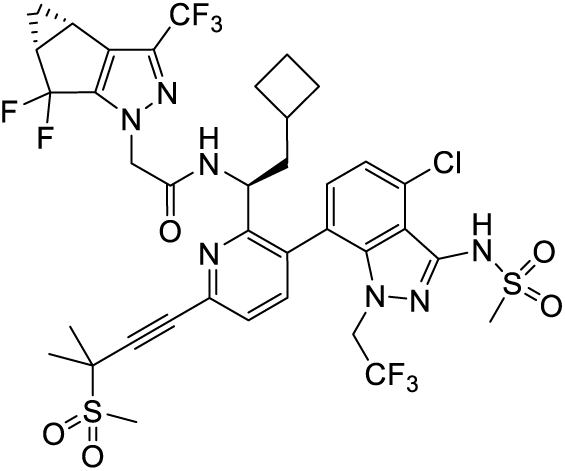

*N-((R)-1-(3-(4-chloro-3-(methylsulfonamido)-1-(2,2,2-trifluoroethyl)-1H-indazol-7-yl)-6-(3-methyl-3-(methylsulfonyl)but-1-yn-1-yl)pyridin-2-yl)-2-cyclobutylethyl)-2-((3bR,4aS)-5,5-difluoro-3-(trifluoromethyl)-3b,4,4a,5-tetrahydro-1H-cyclopropa[3,4]cyclopenta[1,2-c]pyrazol-1-yl)acetamide (**2R**)*.

**^1^H NMR** (400 MHz, CDCl_3_) δ 7.57 (d, *J* = 7.9 Hz, 1H), 7.50 (d, *J* = 5.5 Hz, 1H), 7.48 (d, *J* = 6.3 Hz, 1H), 7.23 (d, *J* = 7.6 Hz, 1H), 7.05 (d, *J* = 7.7 Hz, 1H), 6.65 (d, *J* = 8.8 Hz, 1H), 4.74 (s, 2H), 4.62 – 4.55 (m, 1H), 4.49 (dq, *J* = 16.2, 8.1 Hz, 1H), 3.92 (dq, *J* = 16.0, 8.0 Hz, 1H), 3.39 (s, 3H), 3.14 (s, 3H), 2.52 – 2.41 (m, 2H), 2.06 – 1.94 (m, 1H), 1.81 (s, 6H), 1.75 – 1.58 (m, 5H), 1.57 – 1.48 (m, 1H), 1.45 – 1.38 (m, 1H), 1.36 – 1.29 (m, 1H), 1.17 – 1.06 (m, 1H), 1.04 – 0.88 (m, 1H); **^13^C NMR** (100 MHz, CDCl_3_) δ 164.01, 159.5, 143.5 (t, *J* = 29.3 Hz), 142.6, 139.9, 139.6, 139.0, 136.7 (q, *J* = 39.8 Hz), 132.9 – 132.6 (m), 131.3, 129.7, 127.0, 126.4, 122.7 (q, *J* = 279.9 Hz), 121.7, 120.5 (q, *J* = 267.3 Hz), 119.9 (t, *J* = 243.0 Hz), 119.0, 115.2, 88.2, 85.1, 58.0, 53.7, 50.8 (q, *J* = 34.9 Hz), 49.9, 43.1, 41.5, 35.2, 32.4, 28.3, 28.0 (dd, *J* = 34.3, 29.1 Hz), 27.8, 23.4, 22.7, 22.6, 18.4, 12.3; **^19^F NMR** (376 MHz, CDCl_3_) δ -61.55, -70.33, -81.25 (d, *J* = 256.2 Hz), -103.52 (d, *J* = 255.7 Hz) (minor atropisomer not included); **HRMS** (ESI^+^) Calculated for C_37_H_37_ClF_8_N_7_O_5_S_2_ [M+H]^+^ 910.1858; found 910.1858.

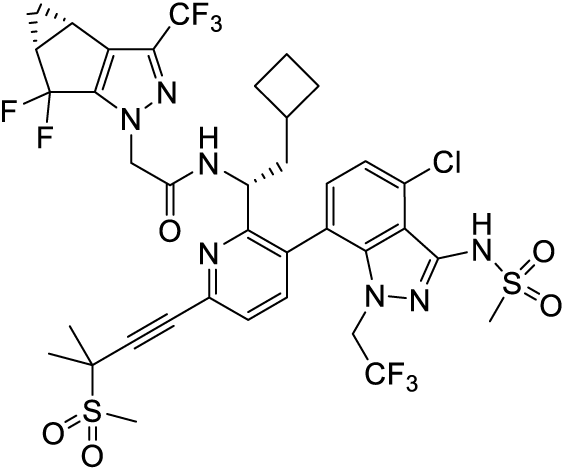

N-((S)-1-(3-(4-chloro-3-(methylsulfonamido)-1-(2,2,2-trifluoroethyl)-1H-indazol-7-yl)-6-(3-methyl-3-(methylsulfonyl)but-1-yn-1-yl)pyridin-2-yl)-2-cyclopentylethyl)-2-((3bS,4aR)-5,5-difluoro-3-(trifluoromethyl)-3b,4,4a,5-tetrahydro-1H-cyclopropa[3,4]cyclopenta[1,2-c]pyrazol-1-yl)acetamide (**3**)

The two atropoisomers of **3** were separated. The NMR of the isolated major atropoisomer is described below, showing a 10% conversion into the minor atropoisomer within hours.

**^1^H-NMR** (400 MHz, CDCl3) δ ppm: 7.57 (d, J = 7.9 Hz, 1H), 7.48 (d, J = 7.9 Hz, 1H), 7.43 (s, 1H), 7.21 (d, J = 7.7 Hz, 1H), 7.01 (d, J = 7.7 Hz, 1H), 6.62 (d, J = 8.6 Hz, 0.9H, major atropoisomer), 6.47 (d, J = 8.9 Hz, 0.1H, minor atropoisomer), 4.75 (d, J = 1.6 Hz, 1.8H, major atropoisomer), 4.72 (d, J = 4.5 Hz, 0.2H, minor atropoisomer), 4.62 (ddd, J = 9.9, 8.5, 3.7 Hz, 1H), 4.53 (dq, J = 16.3, 8.1 Hz, 0.9H, major atropoisomer), 4.41 (dq, J = 15.5, 7.6 Hz, 0.1H, minor atropoisomer), 3.99 (dq, J = 16.1, 8.2 Hz, 1H), 3.41 (s, 3H), 3.15 (s, 3H), 2.54 – 2.43 (m, 2H), 1.82 (s, 6H), 1.63 (ddd, J = 14.1, 9.7, 3.9 Hz, 1H), 1.53 – 1.23 (m, 10H), 1.15 – 1.08 (m, 1H), 0.92 – 0.78 (m, 1H), 0.39 – 0.24 (m, 1H). **^13^C NMR** (100 MHz, CDCl3) δ 164.2, 160.2, 143.7, 142.8, 140.0, 139.7, 139.0, 131.5, 129.7, 127.1, 126.4, 121.7, 119.2, 115.2, 88.1, 85.3, 58.1, 53.9, 51.1, 42.7, 41.7, 36.6, 35.4, 33.0, 31.8, 28.2 (dd, J = 34.4, 29.6 Hz), 25.0, 24.8, 23.5, 22.8, 12.4; **^19^F NMR** (376 MHz, CDCl_3_) δ -61.59, -70.30, - 81.03 (d, *J* = 256.6 Hz), -103.50 (d, *J* = 256.0 Hz). **HRMS** (ESI^+^-TOF) Calculated for C_38_H_39_ClF_8_N_7_O_5_S_2_ [M+H]+ 924.2009; found 924.2024.

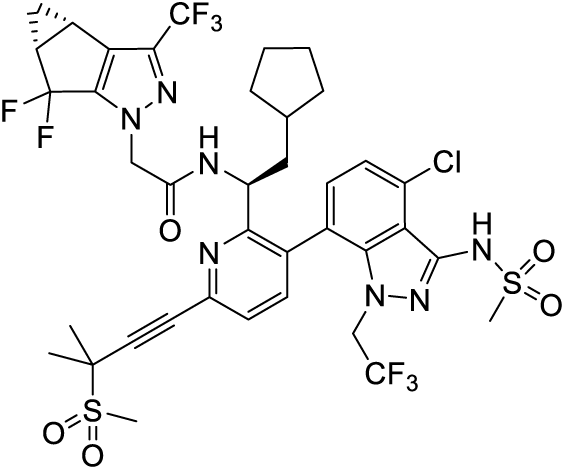

*N-((R)-1-(3-(4-chloro-3-(methylsulfonamido)-1-(2,2,2-trifluoroethyl)-1H-indazol-7-yl)-6-(3-methyl-3-(methylsulfonyl)but-1-yn-1-yl)pyridin-2-yl)-2-cyclopentylethyl)-2-((3bR,4aS)-5,5-difluoro-3-(trifluoromethyl)-3b,4,4a,5-tetrahydro-1H-cyclopropa[3,4]cyclopenta[1,2-c]pyrazol-1-yl)acetamide* (**3R**)

**^1^H NMR** (400 MHz, CDCl_3_) δ 7.60 (d, *J* = 6.9 Hz, 1H), 7.51 (d, *J* = 2.0 Hz, 1H), 7.29 (d, *J* = 1.0 Hz, 1H), 7.22 (d, *J* = 7.6 Hz, 1H), 7.02 (d, *J* = 7.6 Hz, 1H), 6.71 (d, *J* = 8.7 Hz, 1H), 4.77 (d, *J* = 3.3 Hz, 2H), 4.71 – 4.63 (m, 1H), 4.55 (dq, *J* = 16.2, 8.2 Hz, 1H), 3.98 (dq, *J* = 16.0, 8.1 Hz, 1H), 3.41 (s, 3H), 3.16 (s, 3H), 2.53 – 2.41 (m, 2H), 1.82 (s, 6H), 1.69 – 1.58 (m, 1H), 1.55 – 1.47 (m, 2H), 1.46 – 1.39 (m, 3H), 1.37 – 1.29 (m, 4H), 1.17 – 1.09 (m, 1H), 0.92 – 0.78 (m, 1H), 0.36 – 0.24 (m, 1H); **^13^C NMR** (100 MHz, CDCl_3_) δ 164.2, 160.1, 143.5 (t, J = 29.4 Hz), 142.6, 139.8, 139.6, 138.9, 136.7 (q, J = 39.6 Hz), 132.9 – 132.7 (m), 131.4, 129.7, 127.0, 126.4, 122.8 (q, J = 279.6 Hz), 121.6, 120.5 (q, J = 267.3 Hz), 119.9 (t, J = 243.4 Hz), 119.1, 115.2, 88.1, 85.1, 58.0, 53.7, 50.9 (q, J = 34.9 Hz), 50.9, 42.6, 41.6, 36.4, 35.2, 32.9, 31.7, 28.0 (dd, J = 34.7, 29.4 Hz), 24.8, 24.7, 23.3, 22.7, 22.6, 12.3; **^19^F NMR** (376 MHz, CDCl_3_) δ -61.576, -70.30, -81.27 (d, *J* = 256.3 Hz), -103.56 (d, *J* = 256.1 Hz) (minor atropisomer not included); **HRMS** (ESI^+^) Calculated for C_38_H_39_ClF_8_N_7_O_5_S_2_ [M+H]^+^ 924.2009; found 924.2011.

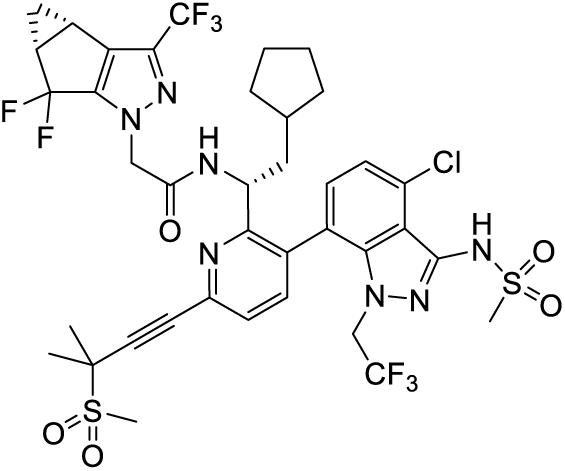

*N-((S)-1-(3-(4-chloro-3-(methylsulfonamido)-1-(2,2,2-trifluoroethyl)-1H-indazol-7-yl)-6-(3-methyl-3-(methylsulfonyl)but-1-yn-1-yl)pyridin-2-yl)-2-cyclohexylethyl)-2-((3bR,4aS)-5,5-difluoro-3-(trifluoromethyl)-3b,4,4a,5-tetrahydro-1H-cyclopropa[3,4]cyclopenta[1,2-c]pyrazol-1-yl)acetamide* (**4**).

The major atropoisomer was separated on preparative HPLC, but it partially converted into the minor atropoisomer, reaching a ratio of ≈90:10 as observed on NMR.

**^1^H-NMR** (400 MHz, CDCl3) δ ppm: 7.60 (d, *J* = 6.3 Hz, 0.1H), 7.57 (d, *J* = 7.9 Hz, 0.9H), 7.48 (d, *J* = 7.9 Hz, 1H), 7.44 (s, 1H), 7.20 (d, *J* = 7.6 Hz, 1H), 7.00 (d, *J* = 7.6 Hz, 1H), 6.59 (d, *J* = 8.4 Hz, 0.9H, major atropoisomer), 6.45 (d, *J* = 9.0 Hz, 0.1H, minor atropoisomer), 4.80 – 4.63 (m, 3H), 4.52 (dq, *J* = 16.2, 8.1 Hz, 0.9H, major atropoisomer), 4.47 – 4.37 (m, 0.1H, minor atropoisomer), 4.00 (dq, *J* = 16.1, 8.0 Hz, 1H), 3.41 (s, 3H), 3.15 (s, 3H), 2.53 – 2.43 (m, 2H), 1.82 (s, 6H), 1.60 – 1.38 (m, 6H), 1.35 – 1.18 (m, 4H), 1.16 – 0.82 (m, 8H), 0.71 (qd, *J* = 12.0, 3.3 Hz, 1H), 0.24 (qd, *J* = 11.8, 3.6 Hz, 1H); **^13^C NMR** (100 MHz, CDCl3) δ 164.3, 160.4, 143.9, 143.6 (t, *J* = 29.4 Hz), 143.4, 142.8, 140.9, 140.0, 139.7, 139.0, 137.03, 136.6, 132.9, 131.5, 129.7, 127.1, 126.4, 124.3, 121.7, 120.0, 119.2, 115.1, 88.1, 85.2, 58.1, 53.8, 51.6, 51.2, 50.9, 49.3, 44.1, 41.7, 35.4, 34.1, 34.0, 31.8, 31.7, 28.2 (dd, *J* = 34.6, 29.2 Hz), 26.3, 26.1, 25.8, 24.8, 23.6, 23.5, 22.8, 22.8, 12.4; **^19^F NMR** (376 MHz, CDCl_3_) δ -61.52 (minor atropoisomer), -61.58 (major atropoisomer), -69.39 (t, *J* = 7.8 Hz, minor atropoisomer), -70.27 (t, *J* = 8.0 Hz, major atropoisomer), -81.05 (dd, *J* = 256.0, 13.4 Hz, major atropoisomer), -81.40 (dd, *J* = 255.5, 13.5 Hz, minor atropoisomer), -103.28 (dd, *J* = 255.9, 9.0 Hz, minor atropoisomer), -103.44 (dd, *J* = 256.1, 8.7 Hz, major atropoisomer); **HRMS** (ESI^+^) Calculated for C_39_H_41_ClF_8_N_7_O_5_S_2_ [M+H]^+^ 938.2166; found 938.2157.

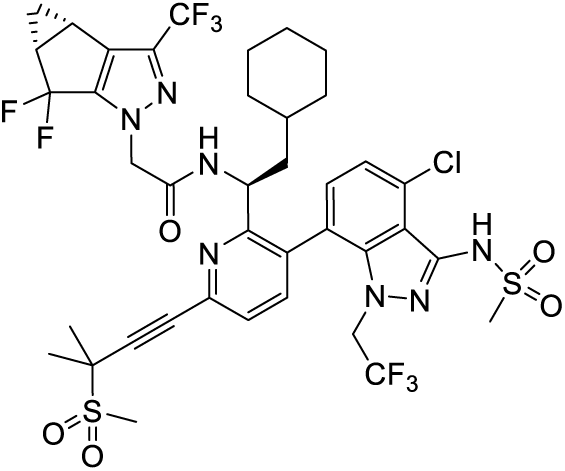

*N-((R)-1-(3-(4-chloro-3-(methylsulfonamido)-1-(2,2,2-trifluoroethyl)-1H-indazol-7-yl)-6-(3-methyl-3-(methylsulfonyl)but-1-yn-1-yl)pyridin-2-yl)-2-cyclohexylethyl)-2-((3bR,4aS)-5,5-difluoro-3-(trifluoromethyl)-3b,4,4a,5-tetrahydro-1H-cyclopropa[3,4]cyclopenta[1,2-c]pyrazol-1-yl)acetamide (**4R**)*.

The major atropoisomer was separated, but it partially converted into the minor atropoisomer, reaching a ratio of ≈90:10 as observed on NMR.

**^1^H-NMR** (400 MHz, CDCl3) δ ppm: 7.57 (d, *J* = 7.9 Hz, 1H), 7.48 (d, *J* = 7.9 Hz, 1H), 7.44 (s, 0.9H, major atropoisomer), 7.29 (d, *J* = 1.4 Hz, 0.1H, minor atropoisomer), 7.21 (d, *J* = 7.7 Hz, 1H), 7.00 (d, *J* = 7.7 Hz, 1H), 4.84 – 4.66 (m, 3H), 4.54 (dq, *J* = 16.3, 8.1 Hz, 0.9H), 4.43 (dq, *J* = 15.0, 7.2 Hz, 0.1H), 3.98 (dq, *J* = 16.0, 8.0 Hz, 1H), 3.41 (s, 3H), 3.15 (s, 3H), 2.53 – 2.42 (m, 2H), 1.82 (s, 6H), 1.60 – 1.38 (m, 5H), 1.35 – 0.82 (m, 9H), 0.71 (qd, *J* = 12.2, 3.3 Hz, 1H), 0.24 (q, *J* = 12.0 Hz, 0.9H), 0.17 – 0.05 (m, 0.1H); **^13^C NMR** (100 MHz, CDCl3) δ 164.2, 160.4, 142.8, 140.0, 139.7, 139.1, 131.5, 129.7, 127.1, 126.4, 121.7, 119.1, 115.1, 88.3, 85.2, 58.1, 53.8, 49.2, 44.2, 41.7, 35.4, 34.0, 34.0, 31.8, 28.2 (dd, *J* = 34.7, 29.4 Hz), 26.3, 26.1, 25.8, 23.5, 22.8, 22.8, 12.4; **^19^F NMR** (376 MHz, CDCl_3_) δ -61.51 (minor atropoisomer), -61.56 (major atropoisomer), -81.11 (dd, *J* = 255.8, 14.0 Hz, minor atropoisomer), -81.17 (dd, *J* = 256.2, 13.5 Hz, major atropoisomer), -103.58 (dd, *J* = 256.4, 8.6 Hz, major atropoisomer), -103.80 (dd, *J* = 256.1, 8.7 Hz, minor atropoisomer); **HRMS** (ESI^+^) Calculated for C_39_H_41_ClF_8_N_7_O_5_S_2_ [M+H]^+^ 938.2166; found 938.8979.

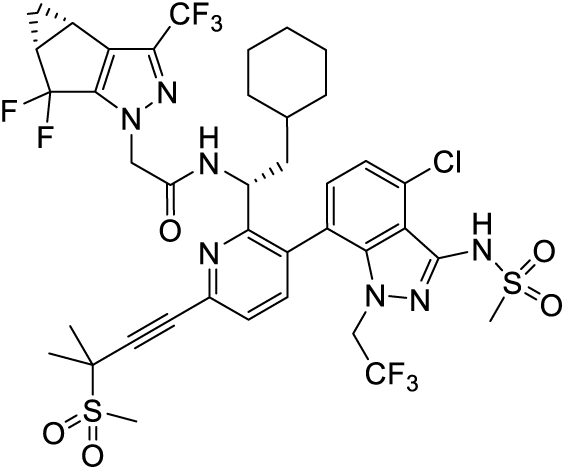

### 4.2. Thermal shift assays (TSAs)

TSAs used purified covalently-crosslinked hexameric CA containing A14C/E45C/W184A/M185A (CA121) either as wild-type (WT) or containing M66I, Q67H, or N74D mutations. CA121 cloned in a pET11a expression plasmid kindly provided by Dr. Owen Pornillos (University of Virginia, Charlottesville, VA, USA). CA121 was expressed in BL21(DE3)RIL *E. coli* and purified according to reported protocols.^42^ The TSAs were performed as previously described,^43-52^ with each reaction containing 7.5 µM WT, M66I, Q67H, or N74D CA121 in 50 mM sodium phosphate buffer (pH 8.0), 1× Sypro Orange Protein Gel Stain (Life Technologies, Carlsbad, CA, USA), and either 1% DMSO (control) or 20 µM compound (1% DMSO final). The plate was heated from 25 to 95 °C with a heating rate of 0.2 °C every 10 s in the QuantStudio 3 Real-Time PCR system (Thermo Fisher Scientific, Waltham, MA, USA). The fluorescence intensity was measured with an Ex range of 475–500 nm and Em range of 520–590 nm. The difference in the melting temperature (ΔT_m_) of CA121 in DMSO (T_0_) verses in the presence of compound (T_m_) were calculated using the following Eq. (1):

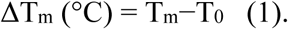

### 4.3. Virus production

The wild-type laboratory HIV-1 strain, HIV-1_NL4-3_,^53^ was produced using a pNL4-3 vector (NIH AIDS Reagent Program, Division of AIDS, NIAID, NIH, Bethesda, MD, USA). The M66I variant was produced by site-directed mutagenesis of the pNL4-3 WT vector in the capsid region. WT or M66I HIV-1_NL4-3_ was generated by transfecting HEK 293FT cells with 10 µg of pNL4-3 vector and FuGENE^®^HD Transfection Reagent (Promega, Madison, WI, USA) in a T75 flask. Supernatant was harvested 48−72 h post-transfection and transferred to MT2 cells for viral propagation. Virus was harvested upon observation of syncytia formation, typically after 3−5 days. The viral supernatant was then concentrated using 8% w/v PEG 8000 overnight at 4 °C, followed by centrifugation for 40 min at 3,500 rpm. The resulting viral-containing pellet was concentrated 10-fold by resuspension in DMEM without FBS and stored at −80 °C.

### 4.4. Anti-HIV-1 assays

Anti-HIV-1 activity of compounds was examined in TZM-GFP cells. The potency of WT and M66I HIV-1 inhibition was determined based on the inhibition of viral LTR-activated GFP expression in the presence of compounds compared to DMSO controls. Briefly, TZM-GFP cells were plated at density of 1×10^4^ cells per well in a 96-well plate. After 24 h, media was replaced with increasing concentrations of compound. Cells were exposed to WT or M66I HIV-1_NL4-3_ (MOI=1) 24 h post treatment. After 48 h incubation, anti-HIV-1 activity was determined by counting the amount of GFP positive cells on a Cytation^TM^ 5 Imaging Reader (BioTek, Winooski, VT, USA) and 50% effective concentration (EC_50_) values were determined. All cell-based assays were conducted in duplicate and in at least two independent experiments. Final values were calculated for each independent assay and average values for all assays were calculated.

### 4.5 Molecular modeling

Two experimental structures of CA were used for molecular modeling: 4XFZ^11^ and 6VKV.^20^ Both structures were prepared using the protein preparation workflow in Schrödinger Maestro (release 2025-3, Schrödinger Inc., New York, USA),^54^ which includes filling of missing hydrogens, side chains, and loops. The resulting hydrogen bonding network was then optimized, and the heavy atoms converged to root mean squared deviation of 0.3 Å using the OPLS4 force field.^55^

Induced fit docking was performed using the induced fit docking protocol using both Glide (Schrödinger Inc., New York, USA)^56^ and Prime (Schrödinger Inc., New York, USA).^57^ The cognate ligand of 4XFZ was selected to be the center of receptor grid in induced fit docking. The ring conformations were sampled using energy window of 2.5 kcal/mol. For docking, receptor (CA) van der Waals scaling was set to be 0.5. Ligand van der Waals scaling was set to be 0.5. The residues within 5 Å of the ligand were refined using Prime.

## Acknowledgments.

This research was supported by the National Institutes of Health (R01AI120860 to SGS and ZW). We thank the Minnesota Supercomputing Institute (MSI) at the University of Minnesota for providing computing resources.

